# Granger-causal inference to study crosstalk between two actin regulators in live cell imaging

**DOI:** 10.64898/2026.11.11.760551

**Authors:** Anteneh Asmare Godana, Tadamoto Isogai, Kushal Bhatt, Meghan Doyal, Gaudenz Danuser, Jungsik Noh

## Abstract

Dynamic actin filament assembly at the cell membrane drives cell migration. Its regulation is governed by functionally overlapping actin regulators and complex feedback loops that hinder conventional perturbation-approaches from dissecting actin regulatory pathways. Here we present an extended, perturbation-free statistical framework based on Granger causality (GC). The framework rests on statistical models that capture the interdependence among subcellular fluctuations of molecular and cell morphodynamic activities extracted from three-channel live-cell imaging. Applied to the inference of cause-effect relationships among actin, its regulators and cell-edge movement, the GC analysis identifies distinct regulatory hierarchies of the actin regulators Arp2/3, VASP and mDia1 on actin dynamics and cell edge movement. Notably, we find that mDia1 plays a critical role in translating actin filament assembly into cell-edge movement, whereas Arp2/3 and VASP function primarily as regulators of polymer extension. By imaging three actin reporters simultaneously, we also demonstrate that the GC framework can distinguish the distinct causal roles of filamentous vs monomeric actin in driving cell edge movements.

## Introduction

Understanding how molecular regulators coordinate to govern cell behavior requires tools that can resolve causality across space and time while avoiding the confounding effects of rapid functional compensation induced by external perturbations, particularly in systems characterized by nonlinear interactions and functional redundancy among regulators. Actin filament assembly during lamellipodia-driven cell migration exemplifies such a system, as it is governed by numerous functionally intertwined actin-associated regulators [1–3]. To overcome the limitations of perturbation-based studies we and others have used spatiotemporal image correlation approaches to establish the temporal ordering of key regulators, including Arp2/3, formins, VASP, and cofilin, relative to actin dynamics, force generation and cell-edge movement [4–10]. However, temporal ordering and correlation alone do not resolve the direction of influence or the direct causal effects between components. Establishing such causal relationships is essential to understand and ultimately engineer molecular regulatory systems characterized by extensive feedback and functional redundancy.

Granger causality (GC), applicable to time series data, statistically determines whether the past dynamics of one variable provides indispensable information for the prediction of another variable’s future dynamics. When applied to live cell time-lapse images, the GC analysis provides a powerful means to identify the directional associations among molecular signals, mechanical events and morphological dynamics, thus enabling useful predictions of cause-effect relationships. To extend GC to spatially structured molecular signals of temporal imaging data, we developed the Spatially Propagating Autoregressive (SPAR) model framework [11]. The SPAR model integrates temporal autoregressive modeling with spatial propagation of subcellular signals to describe subcellular level molecular activities as the cell moves. Thus, it captures information flow between multiple molecular activities across space and time accounting for biochemical continuity, while its multivariate structure controls for potential confounding effects arising from co-localized molecular activities.

Using SPAR models, our prior work reconstructed the causal hierarchy of two cytoskeleton components governing lamellipodial dynamics without any or minimal perturbation. Specifically, we identified a causal chain from Arp2/3 → actin → edge velocity, and further showed that the feedback from actin to Arp2/3 was spatially confined and delineated by the lamellipodium. These analyses also unveiled that VASP and mDia1, but not Arp2/3, exerted additional regulatory effects on cell-edge movement that were distinct from their roles in promoting actin filament assembly. These findings highlighted the potential of causal inference to dissect the multi-functional roles of molecular components within complex regulatory systems.

While the previous SPAR models established the feasibility of inferring regulatory hierarchies from fluorescence fluctuations, they still imposed simplified structures that could not fully capture the multiplexed and redundant regulatory interactions characteristic of actin machinery in the lamellipodium. In this study, we generalize the SPAR framework to four-component causal modeling between three molecules and cell-edge movement. This extension allows us to dissect overlapping actin regulatory pathways and determine how local molecular interactions jointly shape lamellipodial dynamics across subcellular regions. One of the key challenges in this extension is determining the appropriate spatiotemporal lags between components

We designed three progressively refined lag-selection strategies (Approaches 1–3) that balance model complexity with computational efficiency. Because it is computationally prohibitive to evaluate all combinations of time-lags for all possible interactions and spatial propagations among the four components, we adopted a sequential optimization scheme and developed three lag-selection strategies with increasing levels of model complexity. To systematically compare the different strategies, we generated and analyzed 52 three-channel live-cell imaging datasets recording different combinations of Arp2/3, VASP, mDia1 and actin. These datasets also included simultaneous imaging of three distinct actin reporters to quantitatively assess, based on GC measures, which reporter most accurately reflects filamentous actin (F-actin) dynamics and the causal relations between monomeric vs polymeric actin forms and cell edge movement. Together, these examples illustrate the exquisite sensitivity and unprecedented power of GC analyses to unveil causal connections between the constitutive fluctuations of molecular processes that are inaccessible to perturbation-based strategies.

## Results

### Granger causality inference for three-channel imaging data

To develop and validate the four-component GC framework, we repurposed previously acquired three-channel fluorescence time-lapse image sequences of U2OS cells expressing low levels of exogenous mNeonGreen-tagged actin (Actin) and SNAP-tagged VASP (VASP) in a VASP knockdown background, together with endogenously CRISPR-tagged Arp3-Halo (Arp3cr) [11]. Before the present expansion of the GC framework, we were unable to concurrently process the fluctuations of all three molecular components and edge motion, which is necessary to study the interplay between the two regulators Arp2/3 and VASP in the context of actin-mediated cell edge movement. All images were acquired every 3 sec for 301 frames (15 min). We manually cropped the active part of a cell displaying multiple edge protrusion and retraction cycles (Figure 1A). The fluorescent signals of the actin, VASP, and Arp2/3 channels were then sampled in layers of probing windows that maintain a fixed relation to the moving cell edge (Video S1). These signals were transferred timepoint-by-timepoint into image activity maps to visualize their spatiotemporal fluctuations (Figure 1B).

**Figure 1.**
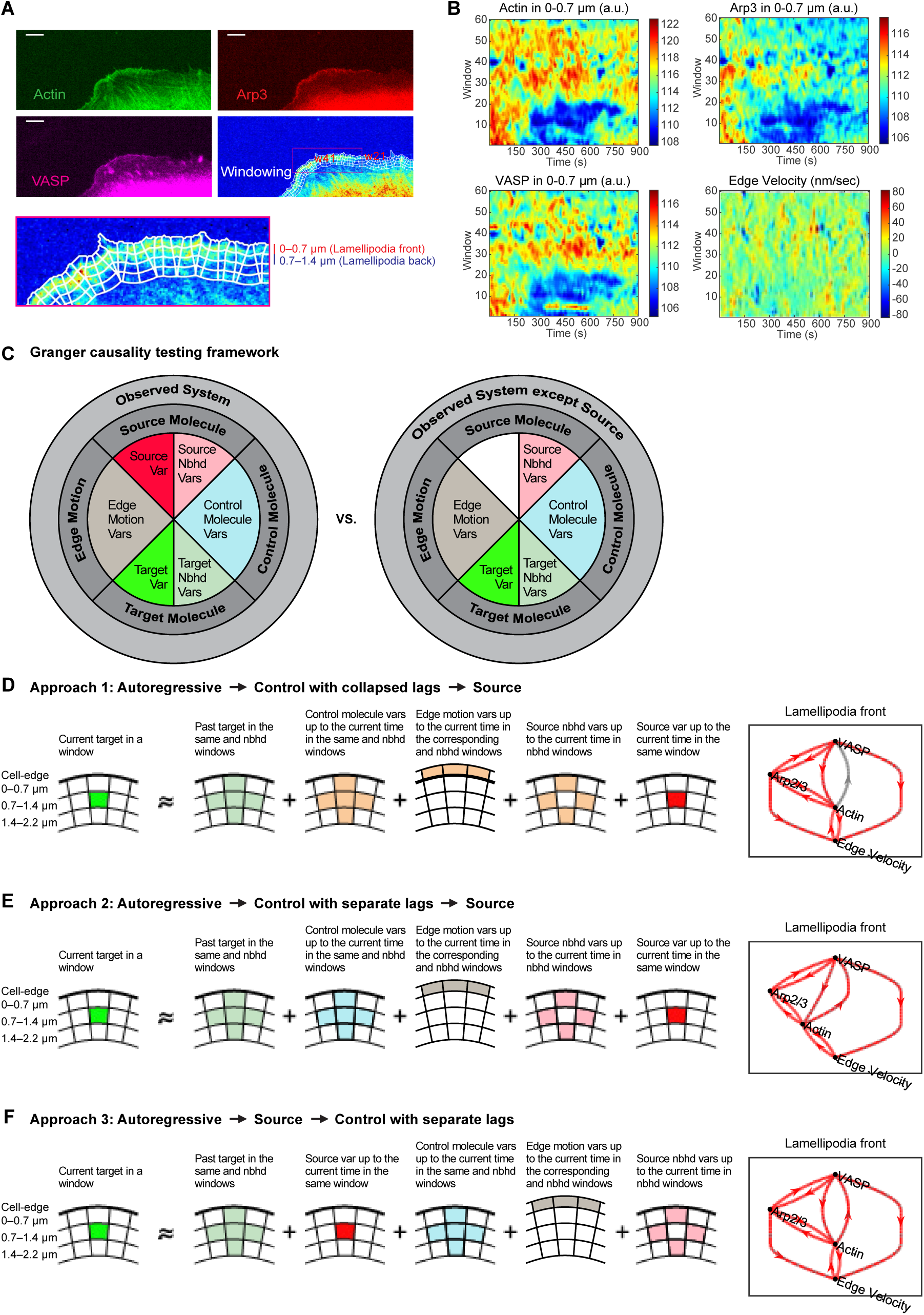
Granger causality framework and model-selection strategies for live-cell cytoskeletal dynamics. (A) Representative fluorescence images of actin, Arp2/3 (Arp3), and VASP at the protruding edge of a U2OS human osteosarcoma cell, illustrating the spatial organization of cytoskeletal components relative to the cell boundary. The lamellipodial front (0–0.7 μm) and back (0.7–1.4 μm) are annotated by concentric spatial windows parallel to the cell edge, which are used for spatially resolved time-series extraction. Scale bars, 5 μm. (B) Representative space–time activity maps of actin, Arp2/3, VASP, and edge velocity within the lamellipodial front, visualizing temporally resolved fluctuations across probing windows. (C) Schematic of the Granger causality (GC) testing framework, illustrating the comparison between two models for the prediction of target molecule activities over time. A full model (left) including the candidate source variable is compared with a reduced model (right) lacking the source variable but retaining autoregressive variables, control molecule variables, edge motion variables, and spatial neighborhood variables. Significantly improved prediction performance of the full model over the reduced indicates a Granger causal connection from the source variable to target variable. (D–F) Schematic representations of the three model-selection strategies used for GC inference. (D) Approach 1 defines the lags of the source variables (red) after optimizing the lags of the autoregressive terms of the target variables (light green) and of control terms (orange). All control terms share one lag structure including source neightborhood variables. (E) Approach 2 uses the same order of lag optimization as Approach 1 but uses separate lag structures for the control terms (cyan, gray, pink), allowing finer separation of conditional contributions to the target signal. (F) Approach 3 defines the lags of the source variables (red) before the lag of the control variables while retaining separate lag structures for the control variables (cyan, gray, pink). The right column visualizes the GC hierarchy between actin, Arp2/3, VASP, and edge velocity for each approach, indicating the substantial impact of different model-selection strategies on the inferred causal hierarchy.

To uncover causal relations among these fluctuations, we developed a GC framework that formalizes the statistical inquiry whether knowledge of the past activity of one component (the source) is indispensable for predicting the future of another component (the target) beyond the contribution from the target’s own past and system covariates [12]. In case indispensability is determined, the source variable is referred to as Granger-causing the target variable. As illustrated in Figure 1C, we structured the system of observed variables into four key categories: the source molecule, the target molecule, the control molecule, and edge motion. To capture spatially adjacent dependencies, each category included variables from the focal probing window as well as from its immediate neighbors. This spatial design is essential for resolving the true direction of information flow in a spatially extended system such as the lamellipodial actin network, where regulatory influences propagate over micron-scale distances (Figure 1B) [13]. The GC test was implemented by comparing a reduced model excluding the source variable at the focal window with a full model including it. Indispensability of the source variable is assessed by the statistical significance of the improvement the full model provides over the reduced model in predicting the target variable. Specifically, in the full model, the current target state in the focal window is predicted by a regression equation simultaneously quantifying the contributions of observed molecular components in focus and neighboring windows and the state edge movement in the associated edge sectors, respectively:

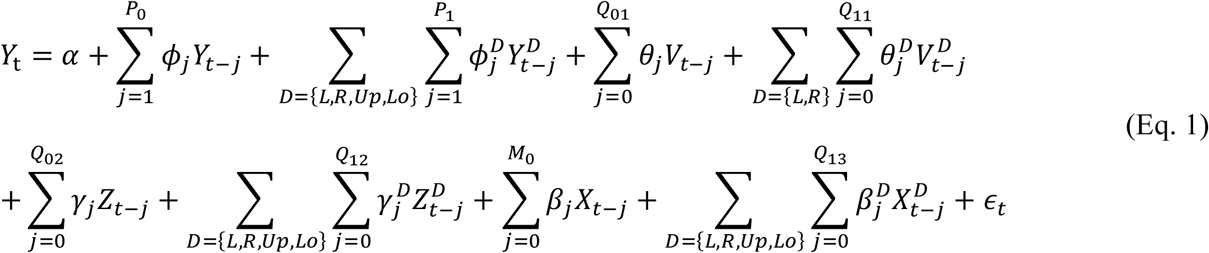

Here, *Y_t_*, *X_t_* and *Z_t_* denote the target, source, and control molecule activities at time *t* ( *t* = 1, 2, …, *T*), respectively. The variable *V_t_* describes the edge velocity. The suffix *D* = {*L*, *R*, *Up*, *Lo*} denotes the left, right, upper, and lower neighboring windows relative to the focus window and *P*_0_, *P*_1_, *Q*_01_, *Q*_11_, *Q*_02_, *Q*_12_, *Q*_03_, *M*_0_ are the temporal lag orders for the different variables. Finally, *ε_t_* describes a random error variable with mean 0 and variance *σ*^2^. The reduced model eliminates the term 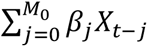 accounting for current and past *M*_0_ source values in the focus window. Importantly, this SPAR model retains the source values in the neighboring windows as potential confounders also in the reduced model. Thus, the GC test strictly determines the indispensability of the source variable in the focus window for the prediction of future values of the target variable in the focus window.

Our previous three-component analysis indicated by an ablation study the importance of including spatial confounders [11]. For example, the model correctly indicated that Arp2/3 was not Granger-causal (G-causal) for edge motion in account of actin, implying that the effect of Arp2/3 activity on edge motion is fully mediated by the actin activity. However, under exclusion of spatial confounders Arp2/3 appeared G-causal for edge motion even in account of actin as a control variable. This false prediction of causality occurs due to indirect information flows that propagate through latent variables, for example, focal Arp2/3 → neighboring Arp2/3 → neighboring actin → neighboring edge motion → focal edge motion. Biophysically, this reflects the fact that actin-regulatory events such as nucleation, elongation, and turnover are not confined to single windows but exhibit short-range coherence due to cytoskeletal connectivity and spatially coupled signaling. Therefore, incorporating spatial context is a necessary condition for inferring causality in systems with both temporal and spatial interdependencies.

### Model selection strategies for Granger causality analysis

To ensure rigorous and interpretable GC inference, we compared three model selection strategies that differ in how they sequentially integrate lagged predictors (Figures 1D-1F). The choice of lag structure, i.e., how many past timepoints to include for each predictor, is critical in biological systems where molecular interactions operate over distinct temporal scales. Without appropriate order selection, models risk either underfitting key dependencies or overfitting noise. Since GC evidence is based on the prediction performances of the full versus a reduced model, each model must be the best predicting model within each class. Thus, accurate lag structure selection is instrumental to GC inference. Exploring all possible lag structures is computationally prohibitive. As a remedy, we designed a strategy in which we sequentially optimized autoregressive lags for the target, lags for control variables including the source neighborhood variables, and the lag for the source (Figure 1D-1F). This approach allows each predictor to contribute at its own biologically relevant timescale, respecting differences in signaling latency and mechanical response. Importantly, we did not adopt the commonly used L1 (LASSO) regularization for model selection because it does not result in interpretable lag structures. In particular, simple L1-regularization assumes implicitly that all predictors influence the system on a similar timescale and apply a uniform sparsity-inducing penalty across different components. This flattens temporal specificity and often obscures weaker but biologically meaningful signals [14].

Our first lag selection strategy (Approach 1) is a straightforward extension of our previous SPAR modeling. We first optimized the autoregressive lag structure for the target variable using data from both the focal and neighboring windows. Next, we introduced control molecule variables, edge motion variables, and source neighborhood signals, all using a shared (collapsed) lag structure. Finally, we introduced the source variable in the focal window with its own individually optimized lag to isolate its unique predictive contribution (Figure 1D). This approach partially separates the source from predictor activity but may still suppress source influence due to shared lag constraints between the source neighborhood variables and other predictors. Approach 2 addresses this limitation by incorporating confounder lag determination in a staged manner. After determining the optimal autoregressive structure of the target, we separately optimized the time-lags for control molecule variables, edge motion dynamics, and source neighborhood activity, followed by the source variable in the focal window with its own lag structure (Figure 1E). While more flexible, this approach can mask source influence if variance is preemptively absorbed by control terms. Thus, in Approach 3 we chose to optimize the source variable lag in advance before optimizing the lags of confounder variables. It first determines the optimal autoregressive structure for the target variable. Subsequently, the source variable is incorporated with its own optimized lag structure, followed in sequence by lag-optimized control molecule variables, edge motion dynamics, and source neighborhood activity (Figure 1F). This design allows each predictor to contribute at its native timescale, enhancing model specificity and improving the interpretability of dynamic causal interactions in the system. In all three approaches the lag structure is determined cell-by-cell using the Akaike information criterion (AIC) as the selection criterion.

Because the computational cost of our extended models increases exponentially with the number of predictors and their independent lag structures, we first performed a systematic lag analysis across the three model-selection strategies (Approaches 1–3) to determine a biologically meaningful upper bound for temporal dependencies. As illustrated in Figure S1, the optimal lag orders inferred from model fits across cells were consistently below 5 frames (15 sec) for all approaches and different relations between actin, Arp2/3, VASP, and edge velocity. This convergence indicates that the dominant causal interactions in lamellipodial dynamics occur within short temporal windows, reflecting rapid feedback among actin assembly, branching, and protrusive motion. Based on this empirical observation, and to balance model fidelity with computational feasibility for large-scale cell populations, we fixed the maximum lag order to 5 for all subsequent analyses. This choice ensured consistent temporal resolution across approaches while substantially reducing computational load in the high-dimensional SPAR based GC framework.

### Approach 2 is more accurate for small sample size cases

We applied the outlined model selection strategies to determine the mutual causal relations between co-expressed mNeonGreen-actin (Actin), SNAP-VASP (VASP), and endogenously Halo-tagged Arp3 (Arp3cr) fluorescent fluctuations and edge movements in U2OS cells [11]. In our previous analysis of this data, restricted to two molecular components, we concluded that Arp2/3 and VASP are independently causal for edge velocity. However, this conclusion had to dismiss the context of actin as the likely central integrator of Arp2/3 and VASP activation.

We compared the G-causal relation networks obtained from the three different approaches after analyzing a relatively small number of live-cell movies (n=6), each harboring between 32 and 70 windows per distance layer. Overall, the different model selection strategies resulted in common G-causal relations between the four components with a few disagreements (Figures 1D-1F). The common GC hierarchy displayed strong actin-centered interactions at the lamellipodia front. Actin significantly predicted future changes in Arp2/3, VASP, and edge motion, underscoring its role as a regulatory nexus for both filament nucleation and elongation machineries and the resulting protrusive dynamics. Likewise, Arp2/3 exerted causal effects on actin and VASP, suggesting a reciprocal feedback between branched actin network formation and actin polymerization. Concurrently, VASP exerted G-causal effects on actin, Arp2/3 and edge motion, consistent with its function as an F-actin polymerase and modulator of branched actin network formation, in addition to its actin polymerization-independent but actin bundling-dependent contributions to edge motion [11, 15–17].

Disagreements between the approaches occurred in the links from Arp2/3 to edge motion and from actin to VASP. We evaluated the three model selection approaches by comparing detailed outcomes of per-cell GC test P-values and the regression lag structures selected from the different approaches, specifically in those conflicting G-causal relations. Our previous work showed strong evidence that the effect of Arp2/3 on edge motion is fully mediated by actin, that is, Arp2/3 is not G-causal for edge motion given actin [11]. Approach 2 reproduced the same causal hierarchy from Arp2/3 to actin and edge motion, while Approaches 1 and 3 predicted actin-independent G-causal effect of Arp2/3 on edge motion (Figures 1D-1F). Interestingly, the examination of per-cell GC test P-values from different approaches showed only slight differences, which however resulted in the opposite GC relations because the per-cell outcomes were at the decision boundary (Figure S1A). We found that notable differences in selected lags, especially between Approaches 1 and 2, led to the different GC outcomes. In Approach 1, assuming a shared lag for all the control variables, the prediction of edge velocity was based on past actin and VASP variables up to the common selected lag of 2 frames (6 sec) (Figure S1A). In Approach 2, optimizing control lags separately, the selected lags for actin and VASP differed. In particular, the selected lag for actin at neighborhood windows was one frame (3 sec) instead of the two chosen in Approach 1, and the lag for neighborhood window VASP was two frames (Figure S1A). Hence, this example demonstrated that the separate optimization of lags for controlling components in Approach 2 improved the accuracy of the prediction model, which could enhance the resolution power in determining G-causal relations especially when the number of movies to analyze is relatively small. Comparing Approaches 2 and 3, the selected lag structures only differed in one cell out of the six cells (Figure S1B). Thus, it suggested that the source prioritized lag selection did not make any improvement and was ineffective in reproducing consistent GC outcomes.

Next, we compared the three approaches in view of their prediction of a causal connection from actin to VASP. Approaches 2 and 3 suggested that the feedback from actin to VASP is significant at the lamellipodia front, while Approach 1 showed weaker GC evidence for the feedback: the rank test P-value was between 5 and 10% (Figure S1B). Again, the more refined model selection in Approaches 2 and 3 resulted in differential lag structures for the control components, where chosen lags were two frames (6 sec) for edge velocity and one frame for Arp2/3 at the focal window in most cells (Figure S1B). Approach 1 selected the common lag of one frame for both edge velocity and Arp2/3 at the focal window. This example again indicated that the refined selection of control variable lag structures enhanced the detection of G-causal effect of the source variable.

Biologically, the causal relationships inferred under Approach 2 delineate a coherent and mechanistically interpretable hierarchy of actin regulation within lamellipodia. At the lamellipodia front, Arp2/3 exerted robust G-causal influence on both actin and VASP, with reciprocal feedback from each, reflecting a tightly coupled module coordinating nucleation and elongation. The resulting actin dynamics were bidirectionally linked with edge motion, both driving and mechanically responding to edge motion dynamics. VASP in parallel exerted an independent unidirectional effect on edge motion, in line with previous analyses [11]. Thus, among the three tested strategies, Approach 2 most effectively resolves both the statistical and biological structure of lamellipodial regulation, yielding a robust and interpretable causal model of dynamic cytoskeletal coordination for a small sample size of n=6 cells.

### Comparison of distinct fluorescent reporters of actin dynamics as indicators of F-actin’s causal influence on morphodynamics

Fluorescently tagged β-actin, which reports both monomeric (G-actin) and filamentous (F-actin) actin dynamics [18], as well as F-actin side-binding peptides such as Tractin that more specifically report F-actin [19], have been widely used to track actin dynamics in cells. Although all actin tracers correlate with edge motion (Figure S2A), they have often been used interchangeably, with no quantitative underpinning as to which tracer most accurately reflects F-actin dynamics. Critically, their highly overlapping localization and dynamic profiles produce strong numerical collinearity in the regression models, making comparison of these three actin reporters an ideal test case for evaluating the sensitivity of the four-component GC framework in delineating functional differentials.

Actin has commonly been tagged at the N-terminus, despite caveats that reporter tags can perturb actin dynamics and disrupt interactions with actin-binding proteins [20–22]. The C-terminus, in turn, is more prone to modifications [23–25] and has been considered less favorable for tagging. To address these uncertainties, we included both N- and C-terminally tagged actin in our comparisons.

Specifically, we endogenously labeled β-actin N-terminally with mNeonGreen (mNG-ACTBcr) and exogenously introduced low levels of C-terminally mScarlet-tagged β-actin (Actin-mScarlet) and SNAP-tagged Tractin (Tractin) in the same cells to minimize cytoskeletal perturbations from co-expression of multiple actin reporters (Figure 2A). Each tracer included a flexible linker of at least 18 amino acids to further reduce structural interference by the tags. We then applied our GC framework to live-cell imaging data from these three actin tracers to assess their causal relationships with edge motion.

**Figure 2.**
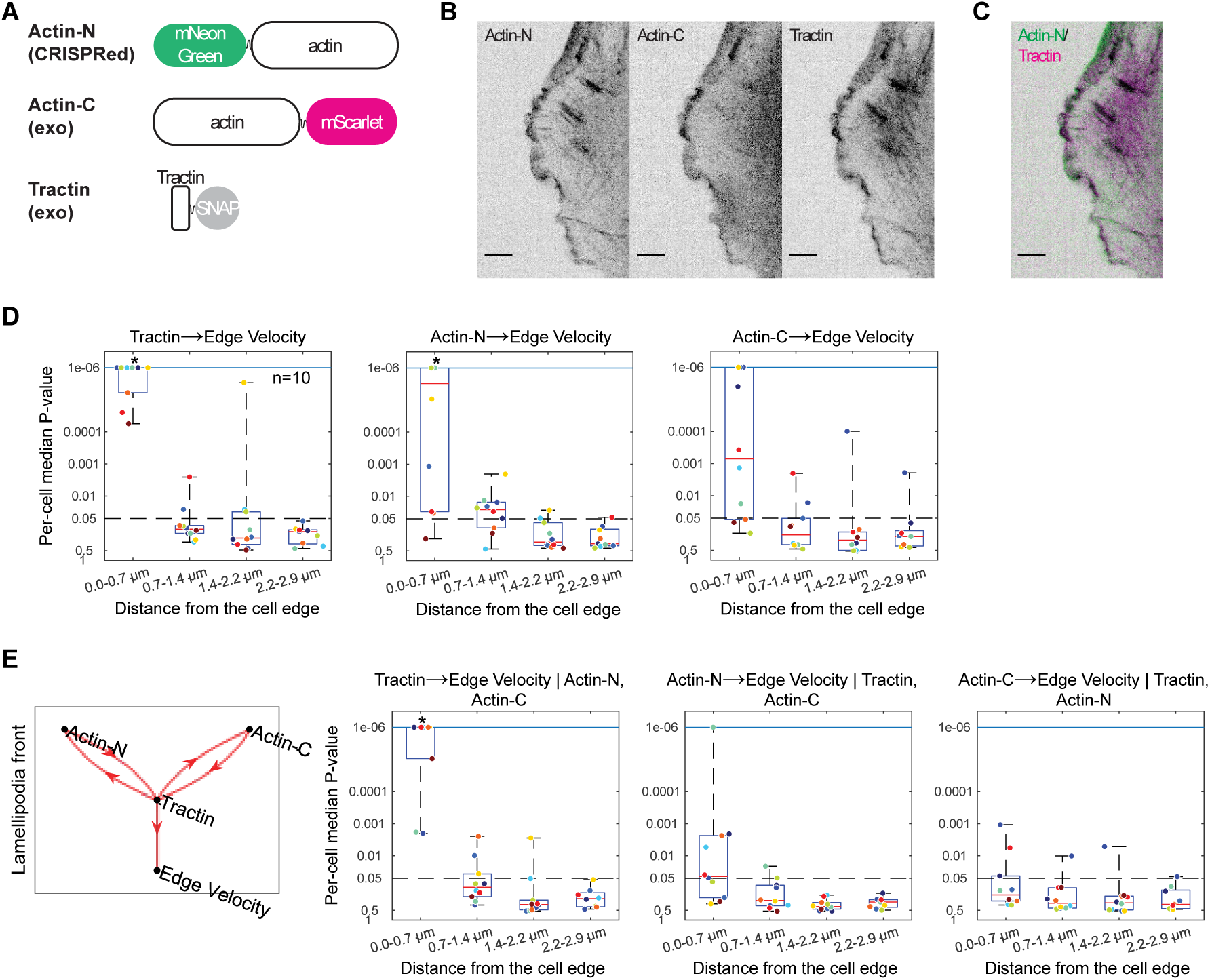
Granger causality inference identifies filamentous-actin is the more direct driver of cell edge motion than monomeric actin. (A) Schematic of actin reporters used in this study, including endogenously tagged Actin-N (CRISPR-edited), exogenously expressed Actin-C, and exogenously expressed Tractin, along with their respective fluorescent labels. (B) Representative fluorescence images of Actin-N, Actin-C, and Tractin near the cell edge, illustrating differences in signal distribution and spatial contrast between the reporters. Scale bars, 5 μm. (C) Actin-N (green) and Tractin (magenta) overlapped. (D) Two-component GC analysis between each actin reporter and edge velocity using Actin-N–based segmentation reveals reporter-dependent differences in actin-edge coupling. Box plots show the distributions of per-cell median GC test P-values at increasing distances from the cell edge (0-2.9 μm, n=10 cells). Data points at a fixed distance denote individual cells, and the dots of the same color represent the same cell. Dashed lines indicate the primary significance threshold (0.05). Asterisks denote statistically significant Granger-causal relations, determined by one-sided Wilcoxon signed-rank tests assessing whether the median of the per-cell median P-value distribution is less than 0.05. (E) Four-component GC analysis by Approach 2. The first column shows the GC hierarchy between the four components. The other columns show the boxplots of per-cell median P-values for the G-causal relations from each actin reporter to edge velocity using the other actin reporters as control components.

Although the three reporters nominally visualize actin, they differ substantially in their labeling of monomeric versus filamentous actin, background fluorescence characteristics, and spatial sharpness of the lamellipodium. These differences, often overlooked in cytoskeletal imaging, proved to have profound consequences for the causal structures recovered specifically within the front-proximal 0–0.7 μm region. Even prior to modeling, raw images highlighted these disparities (Figures 2B and 2C, Videos S2 and S3): Actin-C displayed high cytoplasmic background, frequent intensity spikes, and poorly defined lamellipodial boundaries, whereas Actin-N and Tractin showed clean front-enriched signals with stable lamellipodial geometry. We therefore started our analysis by investigating the stability of the two-component causal relation between edge velocity and each of the three actin reporters separately, first using the Actin-N signal to track the cell boundary (Figure 2D). Causal analysis in a two-component system implies that, for example, the GC from Tractin to edge motion is determined without accounting for Actin-N or Actin-C signals.

Tractin showed highly significant and spatially consistent GC influence on edge motion across all cells (n=10). This matches the expectation that edge motion is directly caused by the polymer turnover of the underlying actin filaments. The distribution of per-cell P-values for the Tractin → edge-velocity relationship clustered tightly in the low ranges, indicating a robust temporal precedence of F-actin assembly over protrusive advancement precisely at the lamellipodia front (Figure 2D). Of note, this causal relation was broken for probing windows at distances > 0.7 μm from the cell edge, which corroborates the previously reported, exquisite sensitivity of the SPAR-model regression to identify causal rather than correlative relations [11]. The Actin-N reporter was also G-causal for edge velocity, however with more modest significance and greater variability between cells (Figure 2D). This behavior reflects the endogenous labeling of both F-actin and G-actin pools. Since Actin-N captures polymerization, depolymerization, and monomer diffusion [20, 21], its temporal fluctuations represent a mixture of protrusion-driving and protrusion-neutral events within the front region, the latter lowering the causal link to edge movement in comparison to Tractin, which reports predominantly the F-actin polymer dynamics. In contrast to both Tractin and Actin-N, Actin-C was rejected as G-causal for edge motion. Its noisy fluorescence, dominated by signals from monomers and most likely also reporter aggregates [23, 25], obscured the fine-scale fluctuations that indicate the causal relation between actin and edge movement. Thus, these GC analyses provide quantitative support to long-standing qualitative arguments in the field regarding the faithfulness of Actin-N and Actin-C as reporters of actin dynamics. Despite concerns that reporter tags perturb actin dynamics and disrupt interactions with actin-binding proteins altogether, N-terminal tagging has been considered the less disrupting approach than C-terminal tagging.

A key reason for this behavior is that Actin-N uniquely combines physiologic biochemical labeling with high-fidelity segmentation geometry at the lamellipodia front. Its endogenous expression preserves the true stoichiometry of actin monomers and filaments, enabling accurate representation of front-localized actin turnover cycles, while its clean spatial enrichment creates reliable boundary definitions for extracting edge trajectories. Unlike Actin-C, whose diffuse fluorescence complicates cell edge detection, Actin-N provides both meaningful molecular information and a geometrically stable template for segmentation. This dual property was crucial for interpreting the segmentation-dependent causal outcomes.

When the cell boundary was segmented from the Tractin signal, we unexpectedly observed that only Actin-N significantly Granger-caused edge velocity, while Tractin and Actin-C did not (Figure S2B). Although Tractin-tracked edge velocities showed small differences to those tracked via Actin-N, Tractin was not G-causal for Tractin-tracked edge movement. It indicated that the reduced diffusive fluorescence in the Tractin signal relative to the Actin-N signal caused the cell edge segmentation to miss critical details in the motion dynamics. Actin-N as a dual reporter of G-actin and F-actin overcomes this limitation, allowing both segmentation and causal prediction at the same time. Actin-C segmentation abolished the two-component GC relations between edge motion and any of the three reporters (Figure S2C). This likely resulted from Actin-C’s low signal-to-noise ratio (SNR) preventing accurate segmentation. Therefore, we continued our analysis using Actin-N as the best-suited fluorescence signal for cell segmentation.

Next, we wondered how the causal relations from two-component GC analysis would translate to a four-component analysis. Remarkably, only the Tractin reporter was significantly G-causal for edge velocity, whereas the Actin-N and Actin-C reporters each had bidirectional G-causal relation with the Tractin reporter (Figure 2E). This indicated that the Tractin signal, as a specific marker of F-actin dynamics contains unique information on the relation between polymer turnover and edge motion not provided by the other reporters. Actin-N was no longer G-causal for edge motion when we account for Tractin together. We also compared the per-cell GC test P-values for the relation Actin-N → edge velocity in two-component analysis (Figure 2D) with those for the relation Actin-N → edge velocity given Tractin in three-component analysis (Figure S2D). It demonstrated that once Tractin was additionally accounted the per-cell P-values became overall non-significant due to the essentiality of the Tractin signal on edge motion. Therefore, the consistently identified essentiality of Tractin over Actin-N that reports both monomeric and filamentous actin dynamics indicates that F-actin is the direct driver of edge motion. This finding emphasizes the sensitive capability of our GC framework to differentiate causal hierarchy between functionally overlapping molecular signals.

The four-component GC hierarchy diagram was supported by all three four-component SPAR modeling approaches (Approaches 1–3). Although the approaches differ in how they introduce lag structures and control variables, the resulting GC network was exactly the same. This demonstrated that when the causality signal is strong and the sample size is large enough, for example n>10, our Approaches 1–3 would be robust and stable determining the same GC hierarchy.

To complement the above discussion of cell segmentation effects on the two-component analysis, we compared the Tractin- and Actin-C-based segmentations to Actin-N-based segmentation in the four-component analysis. Consistently, Tractin was not G-causal for the Tractin-tracked edge movement, while Actin-N was robustly G-causal for the edge velocity calculated from the Tractin-segmentation (Figure S2E). Under Actin-C segmentation, the four-component analysis found no actin reporter is G-causal for the edge velocity as in the two-component analyses. (Figure S2F).

Integrating these findings across all segmentation sources and modeling frameworks yields a coherent mechanistic interpretation of actin regulation in lamellipodia. Tractin is best suited to report F-actin assembly and its functional consequences for cell morphodynamics. However, Tractin is insufficient to demarcate the cell boundary. Accomplishing this necessitates a reporter that backfills the signal from the polymer with a volumetric signal. Thus, both the state of the actin polymer and the space in between the filaments and the edge was achieved by endogenous Actin-N labelling in our case. Actin-N, at least without overexpression, is also suited to faithfully report on the state of F-actin, although its signal is marginally contaminated by the labelled G-actin. This conclusion may change in cell systems or molecular conditions in which the stoichiometry is shifted towards G-actin. Finally, our analyses corroborate the field’s rejection of Actin-C as an appropriate reporter of actin dynamics.

Conceptually, these results underline the exquisite sensitivity of the GC analysis framework to small variations in the fluctuation signals. While the actin reporters define an extreme test case, in which we intentionally enter 2 or 3 nearly identical signals, this sensitivity may translate to causal inference among molecular variables with substantial functional overlap – a key asset of the proposed GC analysis.

### Four-component Granger causality reveals robust VASP-WT feedforward regulation in lamellipodia under VASP-MT interference

Our analysis of three actin reporters demonstrated the exquisite sensitivity of GC analysis in combination with SPAR modeling to detect functional differentials between molecular entities, despite the near-collinearity of variables. In our previous work [11] we had co-expressed a fluorescently labeled wild-type version of the actin elongator VASP (VASP-WT) and a fluorescently labeled mutant VASP (VASP-MT) with attenuated actin polymerization activity in a VASP knockdown U2OS cell line in a control experiment that demonstrated the ability of GC analysis to correctly pinpoint a causal link from VASP-WT to actin assembly while rejecting such a link between VASP-MT and actin. Surprisingly, our analysis predicted that VASP-WT was not only Granger-causal for actin assembly, but that this link was paralleled by a feed-forward causal connection from VASP-WT to cell edge velocity that bypassed F-actin. We interpreted this as an indication of VASP’s dual function as an actin elongator and bundler. Although the wild-type and mutant VASP were co-expressed in the same cell, we were unable to analyze them concurrently as our previous SPAR model only allowed one additional molecule besides F-actin in the analysis. Thus, this data set was an excellent control experiment to test the sensitivity of the newly implemented four-component model in differentiating the functions of wild-type and mutant molecules.

Visually, the characteristics of VASP-WT and VASP-MT are indistinguishable [11]. Indeed, all three numerical approaches (Approaches 1, 2 and 3) reported a strong bidirectional G-causal relation between VASP-WT and VASP-MT (Figure 3A-3C), both at the lamellipodia front and back. While these relations are unlikely to reflect a direct causal interdependence between wild-type and mutant, they indicate that the fluctuations in recruitment of both forms of VASP are modulated by common confounders, such as recruitment to the membrane and cross tetramerization state. Remarkably, despite this tight coupling of VASP-WT and VASP-MT, GC inference in the four-component model still reported functional distinct roles for wild-type and mutant. According to all three numerical approaches, VASP-WT Granger-causes F-actin assembly and independently also edge velocity. As expected, these relations vanish in the second layer of probing windows, at distances > 0.7 µm, demonstrating once again the exquisite sensitivity of GC inference in detecting spatial gradients in the causal connectivity of molecular variables.

**Figure 3.**
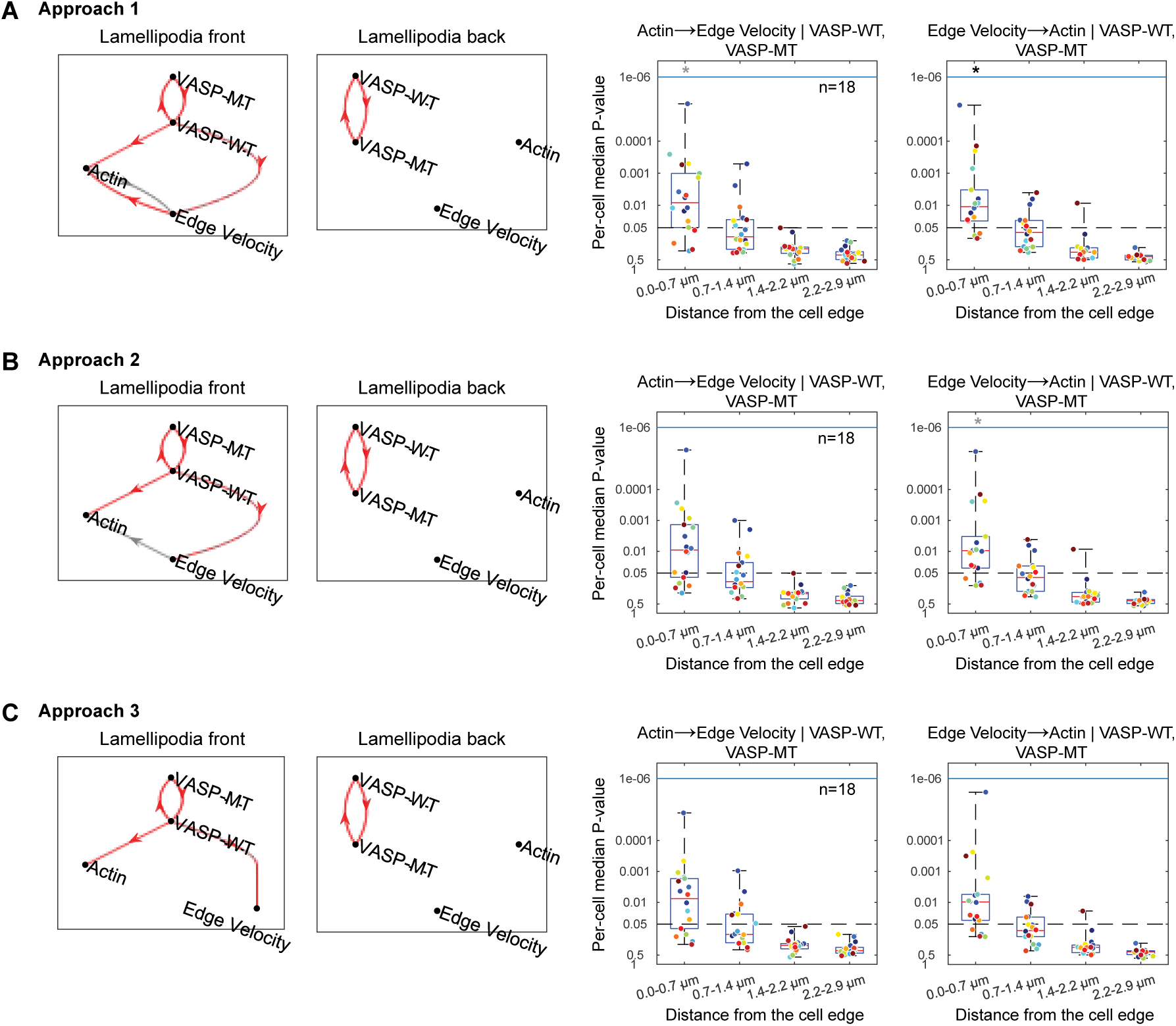
Four-component Granger causal inference confirms VASP’s dual function both as a regulator of actin dynamics and as actin assembly independent regulator of cell edge velocity even when co-imaged with a VASP mutant construct. (A–C) GC hierarchy between VASP-WT, VASP-MT, actin, and edge velocity resulting from four-component analysis using Approaches 1, 2 and 3 (A, B and C, respectively). Hierarchies are shown for the lamellipodial front and back. Solid red edges denote GC relations significant at the standard 0.05 level based on the Wilcoxon signed rank test. Gray edges denote relations significant only at the 0.10 threshold. All three approaches identify consistent causal influence of VASP-WT on both actin and edge velocity despite the presence of the competing VASP-MT signal. VASP-MT has no causal influence on either of these components. However, the three approaches do not detect GC relations between actin and edge velocity due to the multicollinearity with the highly correlated VASP-WT and -MT signals. The two rightmost columns compare the boxplots of per-cell median GC P-values for actin to edge velocity coupling in Approaches 1–3. Black (gray) asterisks denote the significance at the level of 0.05 (0.10).

However, our proposed three modeling approaches differed in determining GC relations between actin and edge motion (Figures 3A-3C). Especially, all three approaches failed to identify the established GC relation of actin → edge velocity at the standard significance level of 5%, which has been strongly identified in almost all imaging data to study the lamellipodia dynamics. At the significance level 5%, only Approach 1 identified the feedback relation of edge velocity → actin (Figure 3A). Additionally at the significance level 10%, Approach 2 detected the feedback edge velocity → actin (Figure 3B), and Approach 1 identified the expected GC relation of actin → edge velocity. Our previous three component analysis of this dataset confirmed the strong GC from actin to edge motion given VASP-WT [11]. Therefore, the failure to detect GC from actin to edge velocity likely stems from the multicollinearity of the VASP-WT and -MT signals, i.e. their time courses were highly correlated. This multicollinearity leads to unstable regression fitting and unreliable model selection [26], ultimately weakening per-cell GC test P-values for the relationship between actin dynamics and edge velocity. Notably, this loss of sensitivity occurred only when VASP-WT and -MT were control components together. Approaches 1–3 identified consistent G-causal relations when VASP-WT or -MT was either of source or target component.

Our GC framework provides a diagnostic measure of multicollinearity for all GC tests at the probing window level (See Methods). We recommend that if the multicollinearity measures across the windows and cells are greater than 30, that is the established threshold, users need to reject the corresponding GC outcomes because the GC tests are inaccurate due to strong multicollinearity. In the regression case of actin → edge velocity given VASP-WT and -MT, the multicollinearity measures were > 30 in a few cells and between 20 and 30 in several cells out of n=18. In contrast, the measures were < 30 for all cells when the regression models did not include both VASP-WT and -MT as control components.

### mDia1 dominates the causal architecture of lamellipodial protrusion, masking actin-edge coupling in four-component models

The extension of SPAR model to four components allowed us to address for the first time how two modulators of F-actin dynamics causally interact with the process of polymer turnover and how this translates into cell edge movement. A particularly interesting case is the interplay between Arp2/3 and the formin family member mDia1. Arp2/3 is widely accepted as the master regulator of branched actin formation in the lamellipodia, determining the rate of F-actin assembly at the leading edge [27, 28], whereas mDia1 is generally associated with the assembly of linear filaments in actin bundles and stress fibers [29, 30]. In a migrating cell the nucleator effect of formins in general and mDia1 in particular is therefore expected to be spatially set back from the protruding edge, co-localized, for example, with focal adhesions where actin bundles grow under the control of complex signaling programs [31]. Indeed, in our three-component GC analysis of Arp2/3 – F-actin – edge velocity vs mDia1 – F-actin – edge velocity, we found consistently strong statistical evidence for the former interactions in the probing windows near the cell edge (0 – 1.4 μm), where the lamellipodia formation occurred. Arp2/3 was non-causal for F-actin assembly in any of the window layers more distal than 1.4 μm, that is, outside lamellipodia [11]. In contrast, mDia1’s nucleator effect emerged in bi-directional causal interactions for the 3 window layers (0.7 – 1.4, 1.5 – 2.2, 2.3 – 2.9 μm).

At the cell population level, the interaction between mDia1 and F-actin was not statistically significant in the front layer of probing windows. However, closer inspection of the distributions of per-cell P-values of GC testing indicated a bimodal distribution. Some cells showed strong G-causal interactions between mDia1 and F-actin at the cell edge whereas other cells showed weak interactions. This behavior seems consistent with the initiation – reinforcement model of F-actin assembly, which predicts that F-actin assembly and cell protrusion is initiated by linear growth of actin filaments under the control of nucleators such as mDia1 and then reinforced by exponential growth of a branched actin network as tension in the expanding membrane rises [6]. Evidence for this model arose; from fluorescent speckle microscopy suggesting the dynamic co-localization of two actin networks at the cell edge with distinct kinetic and kinematic behaviors [4]; from genetic, pharmacological and functional complementation experiments suggesting the requirement of mDia1 for Arp2/3-promoted lamellipodia and ruffle formation [32]; and from fluorescent fluctuation correlation analyses indicating that among a broad set of actin modulators mDia1 is the only molecule recruited to the cell edge significantly before protrusion onset, significantly before Arp2/3 [7]. In a causal analysis without breakdown of phases where the causal structure may change, such as a switch from mDia1 to Arp2/3 as the primary causal driver of F-actin assembly, the outcome of a GC analysis may depend on the fraction of time periods in which the causal relation is active versus inactive. Thus, the bimodal P-value distribution of mDia1 to F-actin causal interaction indicates the coexistence of a subpopulation of movies in which the periods of mDia1-activated F-actin assembly dominated the periods of other nucleator and/or no nucleator activity. While the challenges of detecting transient causal relations remain, especially when they are short-lived compared to the total observation window, we expected the four-component G-causal analysis provide us with much deeper insights into the coordination of mDia1 and Arp2/3 – insights out of reach of any perturbation approach.

To examine the causal interactions between Arp2/3 and mDia1 we exogenously expressed a C-terminally HaloTagged Arp3 (Arp3^EXO^) at low levels in U2OS cells with endogenously tagged mDia1. We first checked whether the three molecular activities causally interact with each other using two-component analyses. If the image fluctuation signals are adequate, any direct or indirect causal linear relationship is supposed to be detected through the two-component analysis. However, we found no G-causal relation between Arp3^EXO^ and F-actin, although Arp3^EXO^ and mDia1 causally interacted with each other (Figure S3A and S3B). In our previous study [11], we had reported a similar observation with mDia1. While exogenous mDia1 in U2OS cells was non-G-causal for F-actin, endogenously CRISPR-tagged mDia1 showed G-causal interactions with F-actin. The GC from endogenous mDia1 to F-actin we observed in the present experiment showed weaker per-cell P-values compared to our previous results (Figure S3C). We suspected that the overexpression of Arp3^EXO^ perturbed the stoichiometry among the nucleators, so that mDia1’s function in the naïve system got masked. Therefore, we generated U2OS cells that stably expressed both endogenously tagged Arp2/3 and mDia1.

With the endogenously tagged Arp2/3 and mDia1, the three different model selection approaches produced near-identical G-causal hierarchy between the four components (Figure 4A). Unexpectedly, the common G-causal hierarchy at the lamellipodia front identified a linear causal chain from Arp2/3 → F-actin → mDia1 → edge motion. Among the three molecules, mDia1 was the closest to edge motion even after accounting for Arp2/3 and F-actin (Figure 4B). F-actin became Granger-non-causal for edge motion after accounting for mDia1, suggesting that mDia1 is a more direct causal factor for edge motion than F-actin (Figure 4B and 4C). Also, mDia1 and Arp2/3 were G-causal for F-actin after accounting for each other, implying that their F-actin regulation follows parallel pathways (Figure 4D and 4E). Statistically, Arp2/3 was G-causal for F-actin only at the significance level of 10%. Given that the GC from Arp2/3 to F-actin has been strongly significant in two-or three-component analyses, this suggests that the number of data points, that is, the length of time courses, might not be fully sufficient to support a four-component analysis with strongly overlapping terms. The only difference between Approaches 1–3 concerned the relation from mDia1 to edge motion. In Approach 1, it was significant only at the level of 10% (Figure 4B). This supports again that Approach 2 is slightly superior to Approach 1 as shown in the above datasets. Thus, we now discuss specific G-causal relations based on Approach 2 outcomes.

**Figure 4.**
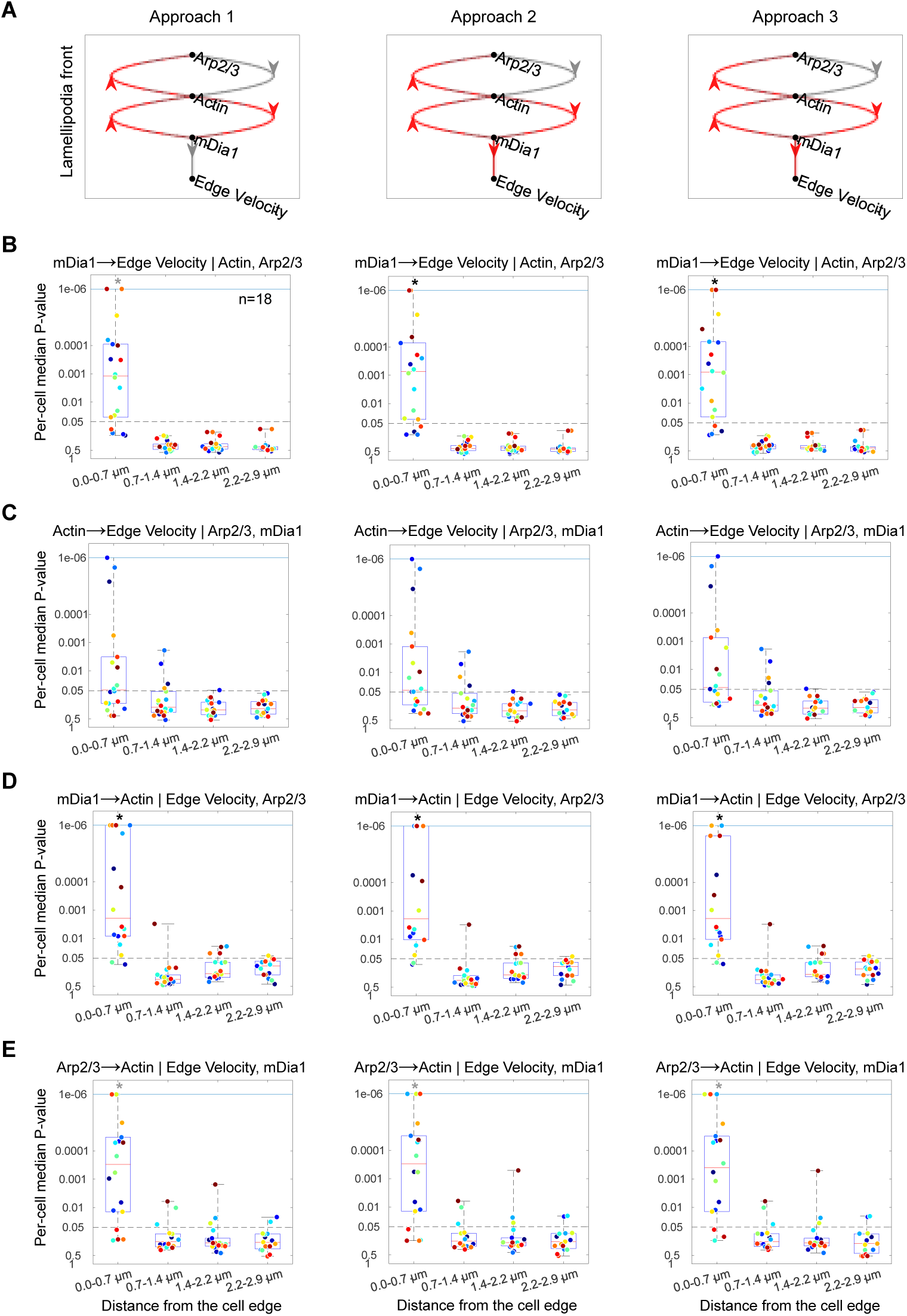
Three different model-selection approaches infer a consistent Granger-causal hierarchy revealing distinct roles of mDia1 and Arp2/3 in actin regulation. (A) Four-component Granger-causal relationship graphs among mDia1, Arp2/3, actin and edge velocity inferred by Approaches 1−3. Red (gray) edges indicate the Granger-causal relationships significant at the level of 5% (10%). Approaches 2 and 3 yield identical hierarchies. Approach 1 results in reduced significance for the Granger-causal connection from mDia1 to edge velocity. (B−E) Comparison of the distributions of per-cell median GC P-values from the different model-selection approaches for key relationships: mDia1 to edge velocity given actin and Arp2/3 (B); actin to edge velocity given Arp2/3 and mDia1 (C); mDia1 to actin given edge velocity and Arp2/3 (D); and Arp2/3 to actin given edge velocity and mDia1 (E). Black (gray) asterisks denote significance at the level of 5% (10%).

Taking a closer look, the endogenously tagged Arp2/3 and mDia1 interacted causally with F-actin in bidirectional fashion (Figure 4A), i.e. both nucleators contribute synergistically, yet causally independent, to polymer assembly at the leading edge. This corroborates the conclusions of our earlier results that two distinctly regulated F-actin populations coexist at the leading edge of migrating cells [4]. Because of the feedbacks from F-actin to both nucleators Arp2/3 and mDia1 are in an indirect causal interdependence. That is, since F-actin is causal for the recruitment of both nucleators, initially mDia1-controlled filaments conceivably are substrates for Arp2/3- mediated branching (mother filament model [32]), whereas mDia1 may be recruited to the barbed-end front of Arp2/3-nucleated filaments. We confirmed this conclusion by identifying that Arp2/3 and mDia1 are indeed G-causally linked in a two-component analysis excluding F-actin fluctuations (Figure 5A). Once F-actin was accounted for in the GC regression model, the per-cell P-values for the GC-relation between Arp2/3 and mDia1 were significantly shifted toward non-causality, showing that Arp2/3 and mDia1 only interact through F-actin assembly and disassembly (Figure 5B).[REF]). Of note, in this particular dataset the per-cell P-value distribution of GC-links from mDia1 to F-actin showed some tendency for bimodality, i.e. some cells lacked systematic causality of mDia1 for F-actin dynamics (Figure 4D). Interestingly, all four cells with a median P-value > 0.05 for Arp2/3 → actin in Approach 2 showed P-values < 0.01 for mDia1 → actin, underlining how these two nucleators compensate for one another in the regulation of F-actin assembly (Figures 4D and 4E).

**Figure 5.**
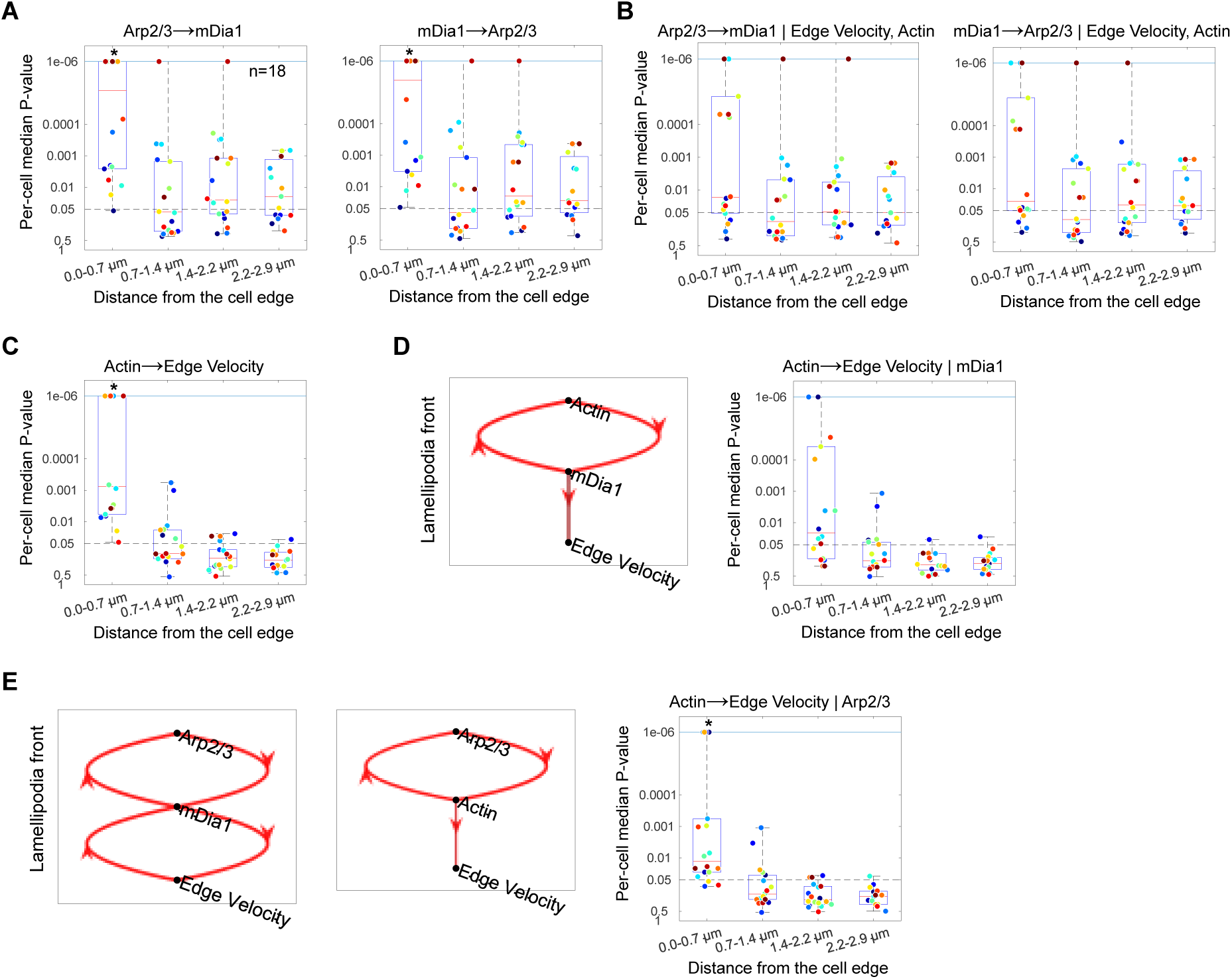
mDia1 indirectly interacts with Arp2/3 through F-actin dynamics and plays a critical role in translating F-actin assembly into cell edge protrusion at the lamellipodia front. (A) Boxplots of per-cell median P-values from two-component GC analysis assessing the GC relation between mDia1 and Arp2/3 without accounting for actin and edge velocity. Black asterisks denote the significance at the 5% threshold (one-sided Wilcoxon signed rank test). (B) Boxplots of per-cell median P-values from four-component GC analysis assessing the GC between the two actin regulators while accounting for actin and edge velocity signals. (C) Boxplots of per-cell median P-values from two-component GC analysis assessing the GC from actin to edge velocity without accounting for mDia1 and Arp2/3. (D) Three-component GC analysis among mDia1, actin and edge velocity. Shown are GC hierarchy (left) and boxplots of per-cell median P-values (right) assessing the GC from actin to edge velocity given mDia1. Actin is not Granger-causal at the lamellipodia front. (E) Three-component GC analyses of the other three-component subsystems, (Arp2/3, mDia1, edge velocity) and (Arp2/3, actin, edge velocity). Shown are the GC hierarchies and the boxplots of per-cell median P-values assessing the GC from actin to edge velocity given Arp2/3.

The most unexpected outcome of the four-component analysis was that, in presence of both nucleators, F-actin was no longer causal for edge velocity, consistently across all Approaches (Figure 4C). Instead, mDia1 was indicated as the sole causal factor for edge movements (Figure 4B). Numerically, this means that fluctuations in the mDia1 fluorescence are more predictive for fluctuations in the velocity time series than F-actin fluctuations. To confirm this critical role of mDia1 on edge movements, we examined related G-causal relations at multiple levels of system complexity through 2-, 3- and 4-component analyses. In a two-component analysis, the per-cell P-values for the GC from F-actin to edge velocity were strongly significant (Figure 5C). Given mDia1, they shifted toward non-causality in the analysis of mDia1 – F-actin – edge velocity, identifying the dominance of mDia1 fluctuation over F-actin in predicting edge velocity (Figure 5D). In addition, the other three-component analyses produced the Granger-causal relationship graphs that were consistent with the result of the four-component analysis (Figure 5E). All those GC analyses of subsystems supported the critical role of mDia1 for edge movements.

Biologically, this does not challenge the notion that F-actin assembly generates the protrusive forces via ratcheted stochastic assembly [33–35]. However, it indicates that at the regulatory level mDia1 plays a central role in how F-actin assembly is translated into cell edge movement. Mechanistically, this role is mediated by mDia1’s function as an elongator of filaments as well as through its competition with other barbed end capping proteins [36, 37]. Moreover, the absence of a G-causal link from F-actin to movement under consideration of mDia1 fluctuations may originate in a portion of actin fluorescence fluctuations that are unrelated to velocity fluctuations. This situation is further complicated by the fact that velocity fluctuations are driven by multiple factors besides F-actin assembly rates, including variation in retrograde flow [4] or membrane-F-actin tethering [38], which the present analysis neglects.

## Discussion

Understanding how cell regulatory circuits with multiple converging target-effector relations and feedbacks operate requires analytical frameworks capable of integrating spatial context between constitutive fluctuations of multivariate molecular activities. This permits the delineation of information flows through the circuit beyond the correlative association between molecular perturbations and phenotype, i.e. shift in circuit behavior. Lamellipodial protrusion is paradigmatic of a cell function governed by such regulatory circuits. The actual protrusion is believed to be downstream of actin filament assembly, which is the nexus for manifold molecular pathways with slightly variable roles. By extending the SPAR-based Granger causality (GC) framework to four-component models, we were able to dissect the directional influences among actin, its nucleators and elongators, and the edge motion output at the leading edge. Three core insights follow from this study: (i) appropriate model selection is required for accurate GC inference especially in small sample size cases; (ii) fluorescent reporter and cell segmentation methods jointly determine whether actin–edge coupling is detectable at the lamellipodia front; and (iii) in a full analysis of actin filament dynamics with nucleators of branched (Arp2/3) and linear (mDia1) polymerization, mDia1 rather than actin emerges as the dominant causal factor for protrusive dynamics.

Expanding our initial SPAR study [11] to incorporate an additional molecular control variable, we tested different strategies for model selection. The selection of regression order, or lags, of the various model components is a most delicate and central step in the implementation of Granger causal analysis. We devised a nested lag-optimization strategy for autoregressive, control and source variables, which preserves the differential temporal scales between molecules and effector variables – here cell edge motion. The convergence of Approaches 1–3 onto nearly identical causal architectures, coupled with the empirical finding that optimal predictive lags consistently fell below five frames for all molecular components, aligns with the sub-minute kinetics of lamellipodial regulation [7, 39]. These results provide strong evidence that the extended SPAR framework is both temporally accurate and methodologically robust.

A second major insight from our work is that the detectability of actin–edge coupling at the lamellipodia front depends critically on both the choice of fluorescent reporter and the fidelity of the cell boundary segmentation. Although actin reporters are often treated as interchangeable, their biochemical properties differ substantially. C-terminal actin fusions have been shown to disrupt actin-binding interactions and alter polymerization behavior [23, 25], while N-terminal tags (Actin-N) preserve stoichiometry and physiologic turnover [20, 21]. Tractin, an F-actin–binding peptide, reliably labels polymerized filaments and provides high spatial contrast at the lamellipodial front [19]. These biochemical differences have profound consequences for GC inference: when segmentation was derived from Actin-N, Tractin provided the strongest G-causal link to edge velocity; Actin-N itself exhibited physiologically meaningful but more variable causality to edge velocity; and Actin-C failed entirely due to its high background fluorescence and unstable signal at the boundary.

Altering the choice of fluorescent probe for edge segmentation indicated the sensitivity of the GC analysis to cell boundary tracking. While Tractin showed a very strong causal link between F-actin and cell edge motion when tracked by Actin-N, the causal relation disappeared when the Tractin signal was also used as the probe for segmentation. The difference between Actin-N and Tractin is that the former contains a diffuse signal phase from monomeric actin reaching the very front of the cell. This signal is absent in Tractin images. Actin-N retained a causal link to Tractin-tracked edge motion, possibly because also the diffusive signal phase is coupled to boundary movements. Under Actin-C segmentation, all causal structure collapsed, reflecting the inability of noisy boundaries to support GC analysis. While Actin-N-based segmentation combined with Tractin provides the most accurate data for the causal analysis of F-actin based morphodynamics, it will generally not be feasible to devote two actin makers to capture this information flow. In practice, imaging Actin-N alone offers the most efficient approach as it generates robust cell-edge tracks and an effective readout of F-actin dynamics.

The four-component analysis allowed us to address widely discuss controversy around the function of Arp2/3 and mDia1. GC analysis supported the hypothesis that F-actin assembly at the leading edge is significantly promoted by the formin family member in addition to the conventional model of an Ap2/3 governed polymer and edge dynamics. This parallelism in nucleator activity is impossible to capture by perturbation of either one of the two nucleators. Importantly, the GC analysis finds no direct causal interaction between the two nucleators. Numerically, this means that the presented SPAR regression framework is sufficiently sensitive to detect differential dynamics in the signal fluctuations of the mDia1 and Arp2/3 probes. Of note, these results rely on expression of both probes at endogenous levels. A slight mismatch in the stoichiometry of the two nucleators disrupts this delicate property of the signals. Biologically, these findings indicate that mDia1 and Arp2/3 contribute differentially to the F-actin turnover, albeit in the same location. They are indirectly causally linked via feedbacks from F-actin, i.e. association of Arp2/3 with F-actin may occur on mDia1-nucleated filaments – referred to as the mother-daughter filament model – and vice versa. How much cross-talk exists between mDia1- and Arp2/3-nucleated filaments cannot be assessed by a nucleator-agnostic probe.

The conceptually most transformative result is the second role of mDia1 as a critical translator of F-actin dynamics into edge movement. This finding challenges the paradigm that the rate of actin polymerization governs edge advancement [40]. Indeed, two-component and partially conditioned three-component models reproduced the expected actin → edge velocity relationship (Figures 5C and 5E). However, when mDia1 and Arp2/3 are included concurrently in the SPAR regression, the causal architecture reorganizes dramatically. In addition to the bidirectional GC interaction with actin, mDia1 exerted a significant feedforward causality to edge velocity (Figure 4). These findings align with previous evidence that mDia1 interaction with F-actin barbed-end contributes to the organization of protrusive actin networks and promotes membrane extension [32, 36, 37]. The dual role of mDia1 as an elongator of actin filaments and as a gatekeeper of the conversion of F-actin assembly into edge movement can be explained by two mutually non-exclusive models. At the level of individual filaments mDia1 competes with capping proteins such as CapZ for association with the barbed end. While mDia1 enables continued growth of filaments and thus forward movement, CapZ association blocks filament elongation. However, this explanation may be too simplistic as CapZ association with some filaments may promote elongation of other uncapped filaments. The proposed GC inference offers an exquisite opportunity to address this complex regulatory circuit between actin filament assembly and cell edge movement. At the level of the entire F-actin network, mDia1’s gatekeeper role may also be explained by a scenario, in which mDia1-driven actin assembly is particularly effective in driving edge movement, whereas the F-actin assembly promoted by other nucleators, including Arp2/3, contributes less to the actual shift in cell boundary. This explanation is consistent with the reinforcement model we proposed before based on direct measurement of F-actin assembly rates by fluorescent speckle microscopy [6, 8]. The reinforcement model projects that F-actin assembly must be highest after the timepoint of fastest edge advancement in order to stabilize the protrusion against increasing membrane tension. Thus, it is plausible that mDia1 causes filament elongation in an early phase of protrusion, in which F-actin assembly directly translates into edge motion, whereas in a later phase Arp2/3 causes growth of branched actin filaments that is, however, no longer fully translated into motion. Thus, both nucleators are causally linked to F-actin dynamics, but only mDia1 association determines when F-actin assembly couples to edge movement.

Collectively, these results underline the power of intervention-free experiments to delineate from constitutive fluctuations of molecular activities the balance of pathways converging on common targets and the multi-functionality of individual nodes in the circuits. They also refine spatially explicit multivariate GC analysis as a generic framework to map out the causal hierarchy in convoluted dynamical systems underlying the regulation of cellular function.

## Resource availability

### Lead contact

Further information and requests for resources and reagents should be directed to and will be fulfilled by the Lead Contact, Jungsik Noh.

### Materials availability

Requests for reagents will be fulfilled by contacting the Lead Contact upon reasonable requests.

### Data and code availability

All original MATLAB code is available at Zenodo (DOI: 10.5281/zenodo.22101054). The code is also available as part of the ‘u-integrate’ umbrella package at GitHub (https://github.com/DanuserLab/u-integrate), which provides tools for live-cell imaging data processing, cross-correlation analysis, and three- and four-component GC analysis.

## Author contributions

Conceptualization: J.N., G.D. T.I.; Methodology: A.G.; Formal analysis: A.G.; Data Curation: A.G.; Validation: A.G., J.N., T.I.; Resources: T.I.; Investigation: G.D., T.I., J.N., A.G.; Supervision: J.N.; Funding acquisition: J.N., G.D.; Writing – original draft: A.G., J.N., T.I., G.D.; Writing – review & editing: A.G., J.N., T.I., G.D.

## Declaration of interests

The authors declare no competing interests.

## Supplemental information

### Supplemental Figure Legends

**Figure S1.**
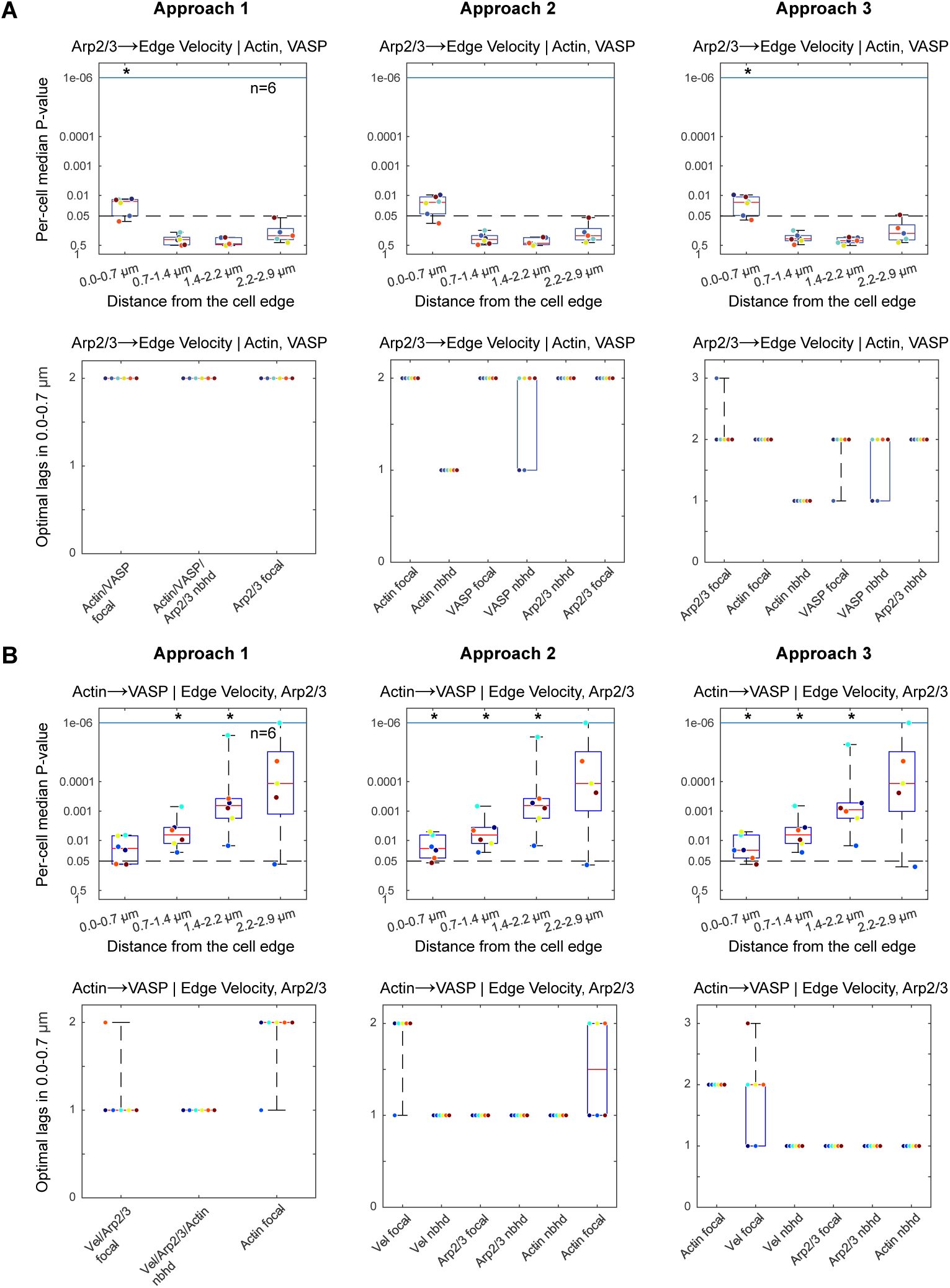
Comparison of different model-selection strategies in the (Arp2/3, VASP, actin, edge velocity) system, related to Figure 1. (A–B) Comparison of three model-selection approaches (Approaches 1–3) across the three columns in terms of per-cell median GC test P-values (top) and selected regression lag structures (bottom). The results are compared in cases when Approaches 1–3 result in contradictory Granger-causal connections: GC from Arp2/3 to edge velocity given actin and VASP (A) and GC from actin to VASP given edge velocity and Arp2/3 (B). Dots of the same color represent the same cell. The asterisks in boxplots of the per-cell median P-values denote statistically significant Granger-causal relationships, determined by one-sided Wilcoxon signed-rank tests.

**Figure S2.**
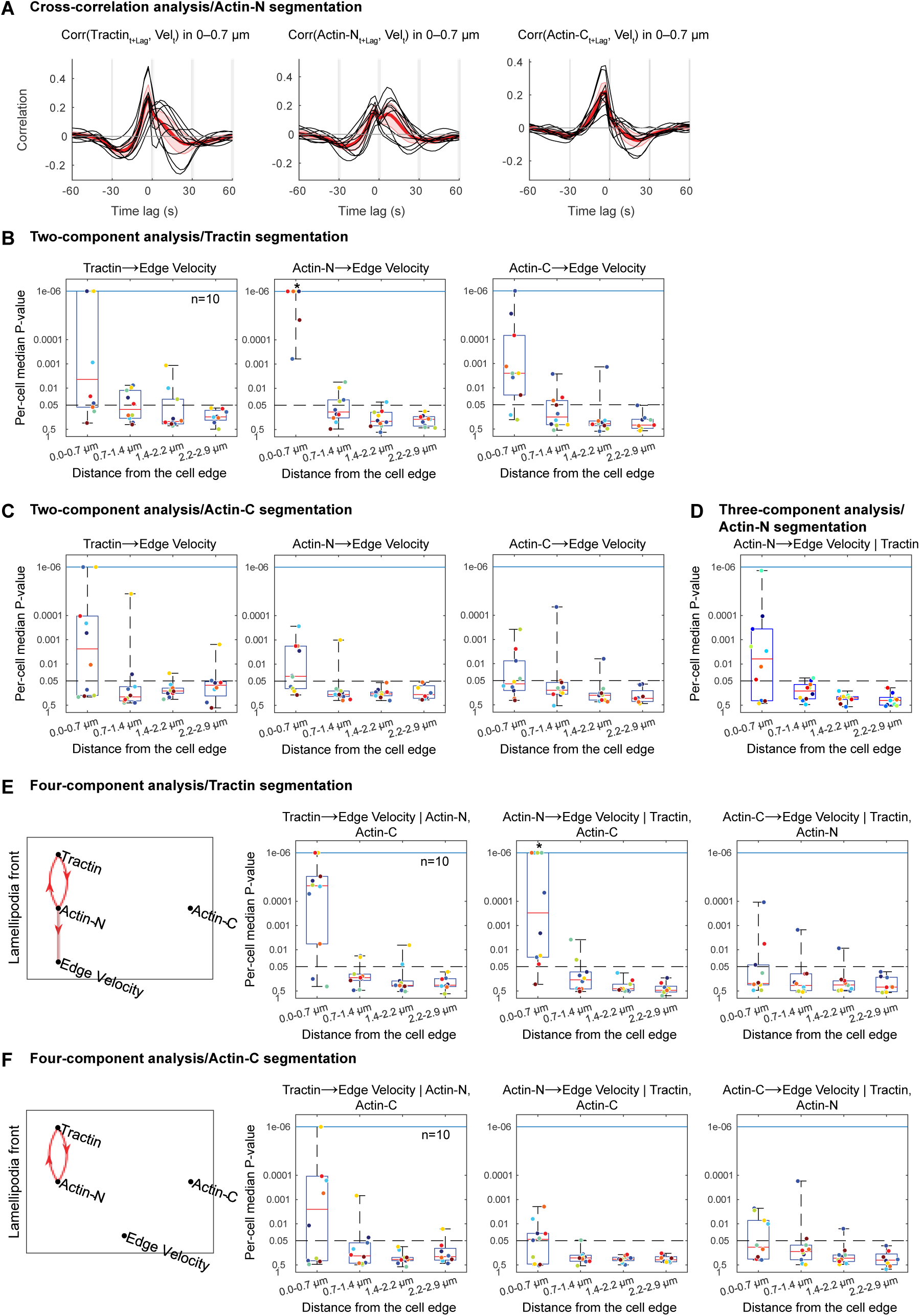
Comparison of distinct fluorescent actin reporters using Granger-causality analysis, related to Figure 2. (A) Cross-correlation profiles between three actin reporters and edge velocity at the lamellipodia front. Cross-correlations between each actin reporter and edge velocity time series are computed at individual probing windows. Black curves represent per-cell averaged cross-correlation curves (n=10). Red curves denote the averages over 10 cells. Shaded bands indicate ± 2 × SEM. The edge velocity is obtained based on the segmentation of Actin-N (CRISPR-edited) images. (B) Two-component Granger-causality (GC) analysis assessing the GC connection from each actin reporter to edge velocity when cell segmentation is based on Tractin images. Boxplots show the distributions of per-cell median GC P-values in 4 probing window layers at increasing distances from the cell edge. Dots of the same color represent the same cell. Asterisks denote significant GC connections, determined by one-sided Wilcoxon signed-rank tests. (C) Two-component GC analysis assessing the same relations as in (B), with cell segmentation based on Actin-C images. (D) Three-component GC analysis with cell segmentation based on Actin-N images. Boxplots show the per-cell median GC P-values assessing the GC from Actin-N to edge velocity given Tractin. (E) Four-component GC analysis with cell segmentation based on Tractin images. The first column shows the GC hierarchy between three actin reporters and edge velocity. The other columns show the boxplots of per-cell median GC P-values from each actin reporter to edge velocity given the other actin reporters. (F) Four-component GC analysis with cell segmentation based on Actin-C images, showing the same analyses as in (E).

**Figure S3.**
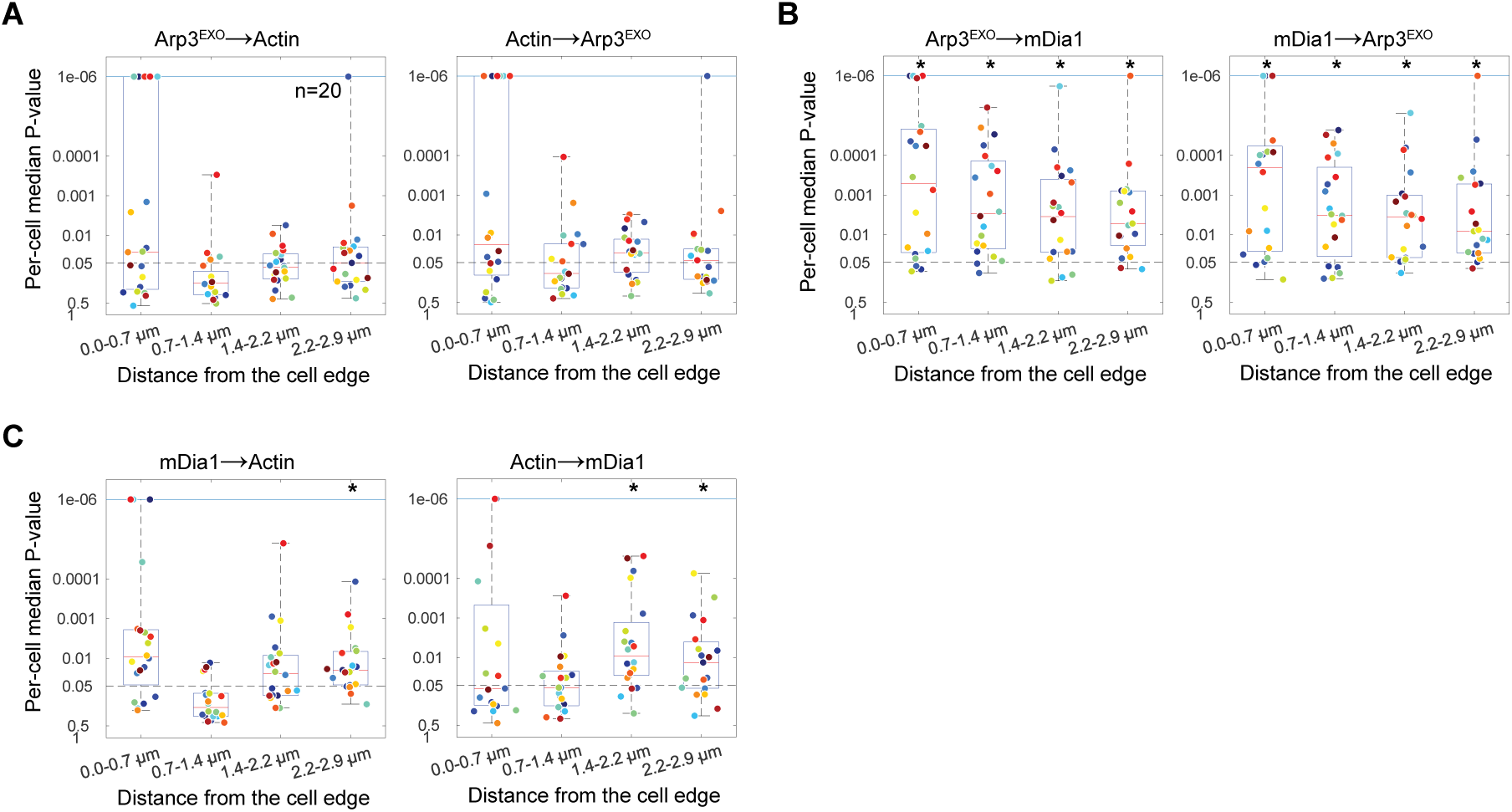
Granger-causality analysis with the imaging of exogenous Arp2/3 and actin and endogenous mDia1, related to Figure 4. (A–C) Boxplots of per-cell median P-values from two-component Granger-causality (GC) analyses based on the time lapse images of U2OS cells expressing exogenous Arp2/3 and actin and endogenously tagged mDia1. The median P-value distributions assess the GC between exogenous Arp2/3 and actin (A), between exogenous Arp2/3 and endogenous mDia1 (B), and between exogenous actin and endogenous mDia1. Asterisks denote significant Granger-causal relationships, determined by one-sided Wilcoxon signed-rank tests with the significance level of 5%.

## EXPERIMENTAL MODEL AND SUBJECT DETAILS

### Cell lines

Human Osteosarcoma U2OS cells were maintained in DMEM supplemented with 10% fetal bovine serum (Sigma; F0926) in a humidified incubator at 37 °C and 5% CO2.

The following U2OS cell lines were previously described [11]: cells expressing exogenous mNeonGreen-actin, SNAP-tagged wild-type and Halo-tagged mutant VASP, cells expressing exogenous mNG21-10 and SNAP-actin in an endogenously mNG211-tandem tagged mDia1 (mNG2-mDia1cr).

β-actin (ACTB) was N-terminally tagged with mNeonGreen at its endogenous loci in U2OS cells (Actin-N cells) by transfecting U2OS cells with pMA-tial1-mNG-18-ACTB donor along with pX330-ACTB-L and R using polyethylenimine (PEI; Sigma). Genome-edited cells were enriched using FACS for mNeonGreen and visually validated under the microscope. C-terminally tagged β-actin (mScarlet; Actin-C) and C-terminally tagged F-Tractin (SNAP; Tractin) transcribed under a truncated CMV promoter to minimally perturb the cells were exogenously expressed in Actin-N cells using lentiviral infection and enriched using FACS.

Cells expressing exogenous Arp3-Halo in mNG2-mDia1cr cells were obtained through infection with lentiviral particles generated from pLVXCMV100-Arp3-HALO, followed by FACS enrichment. Cells expressing both endogenously tagged mDia1 and Arp3 were generated by introducing a C-terminal HaloTag to the endogenous ACTR3 loci of the mNG2-mDia1cr cells with pMA-tial1-ACTR3-18-HALO and pX459-ACTR3. Transfected cells were transiently selected with 1 µg/ml puromycin and single cells sorted into 96 wells plates using FACS.

## METHOD DETAILS

### Plasmids

The following constructs were previously described: pLVXCMV100-mNeonGreen-18-Actin, pLVXCMV100-SNAP-18-actin, pMA-tial1-ACTR3-18-HALO donor vector [11], pLVXCMV100-Arp3-Halo [38]. pSpCas9(BB)-2A-Puro (pX459) and pX330-U6-Chimeric_BB-CBh-hSpCas9 (pX330) were from Addgene (#62988 and #42230, respectively). The guideRNAs previously used to target the ACTR3 loci to endogenously introduce the HaloTag [11] were subcloned into pX459 (pX459-ACTR3). pMA-tial1-mNG-18-ACTB donor vector was cloned by PCR amplifying mNeonGreen flanked with 5’ and 3’ region of ACTB exon 2 surrounding the kozak and start-codon and HiFi assembled into pMA-tial1 [41]. GuideRNAs to target the ACTB loci were cloned into pX330 (pX330-ACTB-L and pX330-ACTB-R). F-tractin-18-SNAP was introduced into the SpeI/MluI-sites of pLVXCMV100 [42] from DNA fragments obtained from a gBlock. β-actin and mScarlet DNA fragments were PCR amplified with corresponding flanking homology arms and HiFi assembled into pLVXCMV100 to generate pLVXCMV100-Actin-18-mScarlet. All primers and gBlocks are listed in Supplementary Table 1. The location and number in the vectors denote the position and the number of amino acids between the tag and the tagged gene, respectively. The 18-linker amino acid sequence was (SGLRSGSGGGSASGGSGS).

### Live-cell imaging of cells

Cells were seeded on fibronectin (10 µg/ml)-coated #1.5 glass-bottom dishes and allowed to spread overnight. The following day, cells were labeled with HaloTag ligand conjugated with JF549 (0.4 – 1 µM; Promega) and/or SNAP ligand conjugated with SiR-647 (0.25 – 1 µM; NEB) for 30 minutes in phenol-red free DMEM supplemented with 20 mM HEPES pH7.4 in the incubator. Cells were then washed twice with phenol-red free DMEM. Time-lapse movies of cells expressing actin, wild-type and mutant VASP were acquired as previously described [11]. All other movies were acquired on a Nikon CSU-W1 spinning-disk confocal microscope equipped with a Hamamatsu ORCA-Fusion sCMOS camera, 100x NA 1.45 oil-immersion objective, Perfect Focus System (PFS), and an Okolab Climate-Controlled Humidified Chamber set at 37 oC. The camera CO2 was turned off because cells were imaged in phenol-red free DMEM supplemented with 20 mM HEPES pH7.4. Each channel was acquired sequentially, with 2×2 binning (130 nm/pixel) and 301 images were acquired every 3 seconds.

## QUANTIFICATION AND STATISTICAL ANALYSIS

### Profiling cell morphodynamics and sampling subcellular fluorescence signals using ‘u-register’

After live-cell imaging, we manually cropped active parts of a cell which display at least a few cycles of protrusion/retraction events by using ImageJ. A set of cropped three-channel videos was first fed into our MATLAB package called ‘u-register’ to segment cell boundaries and track the edge motion together with subcellular probing windows near the edge [13, 43]. Actin channel was utilized for the segmentation unlike our previous study [11], where we used the summation images of the two channel intensities for the segmentation. This choice was motivated by our observation that actin labeled with an N-terminal fluorescent tag is a more accurate marker for the cell boundary than other molecular markers as shown in Figure 2. These image-processing procedures yielded local fluorescence intensity time series as proxies for molecular activities and edge-velocity time series as quantitative measures of the corresponding activity-induced morphological dynamics (Figure 1). Further detailed information on the u-register MATLAB package is available in a Danuser lab GitHub repository (https://github.com/DanuserLab/u-register).

### Preprocessing steps for Granger causality (GC) analysis

With the obtained spatiotemporal readouts of molecular activity and edge motion velocity at the subcellular level, the following preprocessing steps were implemented to ensure that input time series data satisfied the statistical assumptions required for the subsequent regression analysis and GC tests. First, we filtered out quiescent edge segments before inferring GC relations, because we have observed no significant correlative and causal relations between the actin, its regulators and edge velocity within the quiescent edge segments. We implemented Ljung-Box test [44] using a MATLAB function *lbqtest()* to test if an edge velocity time series is a white noise, which implied that the corresponding probing window showed no systematic protrusion/retraction dynamics. Second, we subtracted out low-frequency oscillations from all the molecular activity time series. We observed that the average duration of protrusion/retraction cycles was maintained at ∼60 sec in our U2OS cell system. But the involved molecular activity fluctuations typically exhibited longer cycles. Thus, we applied a simple normalization to the molecular time series to subtract out the oscillation components slower than the average protrusion/retraction cycle of ∼60 sec. This low-frequency subtraction (LFS) was designed to mainly improve the cross-correlation curve output between molecular activity and edge velocity. Because applying the LFS resulted in no substantial difference in the identified GC relations as reported previously (Noh2022), users may skip the LFS step. Third, we checked whether each edge velocity or molecular activity time series displayed a deterministic trend or not and excluded a video if more than half of its probing windows contained such non-stationary time series from the GC inference. Because the GC inference requires given time series to have no deterministic trend, that is, the input time series need to be statistically stationary, we implemented an augmented Dickey-Fuller test [45] to automatically identify the fluctuation signals with a trend. Further details of the preprocessing steps can be found in our previous study [11].

### GC inference using ‘u-integrate’

We previously developed and deployed a MATLAB package called ‘u-integrate’ to perform GC inference with two channel fluorescence imaging data. Through this study, we augmented three new pipelines corresponding to Approaches 1–3 for three channel imaging data, which are available at a public repository (https://github.com/DanuserLab/u-integrate). The u-integrate pipeline started with the preprocessing steps described above, and then the full and reduced regression models were fitted at each subcellular window after exhaustive AIC-based model selection. For each candidate directional relation, the GC P-values were computed at individual windows using F-test that compares the full and reduced models. These window-level P-values were then summarized for each cell by their median. The resulting per-cell median P-values from *n* independent cells were used to assess GC relationships between actin and its regulators near the cell edge, as visualized in Figure 2.

To determine whether a GC relationship was consistently supported across cells, we applied a one-sided Wilcoxon signed-rank test to evaluate whether the distribution of per-cell median P-values was significantly shifted below the nominal significance level of 0.05. A significant result indicated that the typical per-cell median P-value was less than 0.05, implying that, more than half of probing windows within the corresponding layer in a cell provided evidence supporting the proposed GC relationship. As the final output, the u-integrate pipeline generated layer-specific GC network diagrams summarizing the inferred directional interactions among the three imaged molecular components and edge motion. In addition, the pipeline produced a comprehensive set of visualization and diagnostic outputs for interpreting and validating the regression analyses, including residual plots, fitted regression coefficient profiles, selected model orders, AIC curves, and other model diagnostic summaries. Further details on the F-test and regression equations were described previously [11].

### Statistical diagnostics of GC models in individual probing windows

To ensure the validity of GC inference at the single-window level, we evaluated each fitted regression model using three diagnostic criteria: residual autocorrelation, multicollinearity, and the normality of residual distributions. All regression analyses reported in this study satisfied these diagnostic criteria, except the multicollinearity in actin, VASP-WT and VASP-MT dataset as described below.

First, autocorrelation functions (ACFs) of residuals were examined for each fitted model. Valid GC inference requires residual autocorrelations to be close to zero across all lags, indicating that the selected lag structure adequately captured the temporal dependence in the data.

Second, multicollinearity among predictors was evaluated using the condition number of the regression design matrix. The condition number provides a global measure of collinearity among regressors, with large values indicating near-linear dependence and unstable parameter estimates [26]. Models with condition numbers exceeding 30 are considered severely ill-conditioned. Only in actin, VASP-WT and VASP-MT dataset, we observed that a few cells displayed many ill-conditioned regression cases with the condition number > 30, and that several cells showed high multicollinearity cases with the condition number between 20 and 30.

Third, residual normality was evaluated using the Kolmogorov–Smirnov (KS) test. Though residual normality was well-satisfied in almost all probing windows and regression cases in this study, it is worth noting that the GC inference is relatively robust to mild departure from the normality of residual distributions.

## Notes

### Competing Interest Statement

The authors have declared no competing interest.

